# Limited genetic structure and high gene flow in *Fasciola hepatica* populations infecting ruminants in different geographic areas in the UK

**DOI:** 10.64898/2026.04.01.715781

**Authors:** Muhammad Abbas, Kezia Kozel, Nick Selemetas, Olukayode Daramola, Eric R. Morgan, Umer Chaudhry, Martha Betson

**Author notes:** Corresponding author: Martha Betson.

## Abstract

The liver fluke, *Fasciola hepatica*, is a major parasitic threat to ruminant health and productivity worldwide, with important implications for food security, animal welfare, and zoonotic risk. This study developed and validated a multiplex deep amplicon sequencing assay targeting the mitochondrial NADH dehydrogenase 1 (mt-ND1) and cytochrome c oxidase subunit 1 (mt-COX1) loci for high-throughput genotyping of *F. hepatica*. DNA was extracted from eggs sedimented from sheep and cattle faeces (n = 78) received from farms and from adult worm pools (n = 12) isolated at abattoirs from diverse regions across the UK. Following high-throughput sequencing, bioinformatics analysis was performed to demultiplex Illumina sequence reads and extract amplicon sequence variants (ASVs). A total of 11 ASVs were identified at each locus (mt-ND1: 264–279 bp; mt-COX1: 312–319 bp), with two or three predominant ASVs per locus, along with rare variants. Network and PCA analyses revealed two distinct clusters at the mt-ND1 locus: one primarily associated with sheep and another shared between sheep and cattle. In contrast, mt-COX1 sequence reads formed a single dominant cluster. Population analyses revealed extensive ASV sharing across regions, indicating high gene flow, likely facilitated by livestock movement and parasite adaptation.

## Introduction

The liver fluke genus *Fasciola* comprises two species: *Fasciola gigantica* and *Fasciola hepatica*. *F. hepatica* is prevalent in temperate zones, including the UK, Europe, parts of Oceania and the Americas [1], however, both species can be found in tropical and subtropical regions of Asia and Africa, and their hybrids are also prevalent in Asia and Africa [1,2]. The lifecycle of *Fasciola* is complex, with various definitive mammalian hosts, including sheep and cattle. The intermediate hosts in this lifecycle are mud snails: *Galba truncatula* [3], previously known as *Lymnaea truncatula,* for *F. hepatica* [4] and *L. natalensis* for *F. gigantica* [5]. This parasite is transmitted through food plants and herbage contaminated with metacercariae, infecting mainly small and large ruminants, though other mammalian species including humans can be infected [6].

Various factors can influence the occurrence of the parasite and resulting disease, including environmental conditions such as rainfall [7], moisture levels, and temperature (with temperatures from 10°C to 25°C being optimal), as well as the geography of grazing areas (e.g., topography and soil type) [8–11] and animal movement [12]. A feature of the lifecycle is the clonal expansion of *Fasciola* spp. within its intermediate snail host, which contributes to pasture contamination and to the subsequent infection of hosts by metacercariae of the same genetic origin. This clonal expansion may lead to a genetic bottleneck effect in the parasite, particularly when infection levels in snail populations are low [13].

Understanding the population genetics of *F. hepatica* infection provides crucial insights to aid the design of effective control strategies [14]. Over the past few decades, high levels of animal movement have been reported in domestic ruminants in several European countries [15,16]; hence, analysing the population genetic structure of *F. hepatica* can assist in understanding the corresponding spread of parasites infections. Determining parasite population genetics can further inform on transmission dynamics and infection rates, thereby helping identify interventions to reduce disease burden [12]. For example, if a parasite population is characterised by a single dominant amplicon sequence variant (ASV) at high frequency, this suggests a single infection source with clonal expansion of the parasite in the intermediate host, with low metacercariae mixing in pasture settings [12,13]. On the other hand, multiple ASVs at varying frequencies in parasite populations might indicate multiple infection sources on the farm and high mixing of metacercariae [13,17]. Few studies of large and diverse fluke populations examine whether infection has emerged recently in the host at a single time point, or whether burdens have been established repeatedly at different times before spreading. Recently, we have used these methods to study the multiplicity of *Calicophoron daubneyi* infection in the United Kingdom [17] and *Fasciola gigantica* infection in Pakistan [12]. Our findings were consistent with multiple independent emergences of *C. daubneyi* infection, while the identification of common variants across several populations spanning a range of geographic locations highlights the role of animal movements in the parasite’s spread [17]. Moreover, our findings also suggest that most of the hosts were predominantly infected with the emergence of *F. gigantica* infection, while the identification of identical variant, consistent with clonal multiplication within the snails. The most common variants was identified across several populations spanning a range of geographic locations, again highlighting the role of animal movements in the spread of *F. gigantica* infections [12].

Population genetics can be determined using mitochondrial DNA (mtDNA) markers [18–22], as well as nuclear microsatellite loci [23]. Microsatellites are usually highly polymorphic 1-6 bp sequences that can be used as markers to investigate genetic diversity and genetic differentiation using genomic DNA [24]. Recently, a panel of 15 highly polymorphic nuclear microsatellite loci tested on DNA extracted from different life cycle stages, such as eggs, adult worm, miracidia and metacercariae has been reported for *F. hepatica* [23]. High levels of genetic diversity and clonal expansion of parasite and panmictic (randomly mating) populations has been reported by using microsatellite markers on an abattoir-based population genetics study of *F. hepatica* in cattle in the central England and Wales [13]. There are several advantages of using microsatellite markers include their distribution over eukaryotic nuclear genomes [25], and mutation information can be useful for calculating Hardy-Weinberg Equilibrium (HWE) to study homozygosity and heterozygosity [23,24] because genomic DNA contains information on inheritance from both parents [26]. However, limitations include high mutation rates and elevated levels of polymorphism [27]. Furthermore, a high number of alleles per locus in microsatellites can inflate F-statistic values [28] and confound interpretation. Thus, this issue can sometimes lead to over- or underestimates of genetic diversity when the most common allele occurs at either very low or very high frequencies [29]. Microsatellite datasets can be also prone to genotyping errors, which can bias downstream population genetic analyses [30].

Mitochondrial DNA is frequently used as a marker to study population genetics because of its haploid nature [31], maternal inheritance [32], high copy number [33], clock-like and neutral evolution rates [34], and a lack of recombination after heteroplasmy [35]. These characteristics also make mitochondrial markers valuable for investigating the genetic diversity and genetic differentiation in fluke populations. Mitochondrial markers have been well documented for studying population genetics in *Fasciola* spp. from different regions of the world [12,14,22].

As sequencing technology advances, deep amplicon sequencing enables the study of the population genetics of *Fasciola* spp. in greater depth [12,21,23]. For population genetics study,the mitochondrial marker mt-ND1 was used for *F. gigantica* and mt-COX1 for *C. daubneyi*, along with deep sequencing, utilising DNA isolated from adult worms of naturally infected cattle and sheep [17,36]. Deep sequencing enables the detection of dominant and low-frequency variants in different parasite populations [36], which can be useful to study insights into parasite transmission intensity, infection sources, and metacercarie mixing. However, assays for assessing genetic diversity of *F. hepatica* using deep sequencing of multiplexed mitochondrial markers yet not exist. Such assays would provide opportunities to shed new light on the population genetics of *F. hepatica* from naturally infected hosts across various regions of the UK. For example, the extent of overlap between sheep and cattle, and the presence of geographical clustering, could provide insights into the roles of pasture sharing and livestock movement in driving fluke infection and disease.

The present study aimed to develop and test an improved deep amplicon sequencing approach to investigate the population genetics of *F. hepatica* infections across the UK using two mitochondrial markers (mt-ND1 and mt-COX1). This multiplexed sequencing method and high-throughput sequencing approach reduces experimental complexity for studying host-level population genetics in *F. hepatica* populations using a single Illumina sequencing run. This methodology enabled the examination of transmission dynamics and gene flow in both adult worm and egg DNA obtained from natural *F. hepatica* infections in the UK.

## Results

### Validation of multiplex PCR and demultiplexing of mitochondrial markers

The multiplex meta-barcoded PCR targeting mitochondrial positive control *F. hepatica* DNA successfully amplified distinct bands at 311 bp for mt-ND1 and 359 bp for mt-COX1, with no evidence of non-specific amplification (Supplementary Fig. 1a, b and c).

Out of a total of 90 individual samples processed, 78 (86.66%) samples generated sequence reads for the mt-ND1 marker (n = 12 adult worm DNA, n = 66 egg DNA). These sequences obtained from mt-ND1 loci were categorised into 40 parasite populations, comprising cattle (n = 14) and sheep (n = 26) populations. For the mt-COX1 marker, 84 (93.33%) samples successfully produced sequence reads (n = 12 adult worms, n = 72 egg DNA), grouped into 42 parasite populations, including cattle (n = 15) and sheep (n = 27).

### Geographical distribution of mt-ND1 locus ASVs of *F. hepatica*

Eleven ASVs were identified at the *F. hepatica* mt-ND1 locus (accession numbers PX902280-PX902290), ranging from 264 bp to 270 bp, using a library of mt-ND1 reference sequences downloaded from NCBI database (Supplementary Fig. 2), and their frequencies were recorded across 40 fluke populations in different counties of the UK (Fig. 1a and Supplementary Table 1). Across the 40 parasite populations analysed, a total of 1,403,462 sequence reads were extracted, of which the majority (1,167,674 reads; 83.1%) belonged to a small number of predominant ASVs, including ASV1, ASV2, and ASV3. These predominant ASVs reflect the dominant variants circulating in cattle and sheep in different regions. ASV1 was the most abundant variant (39% of total reads), detected in 18 populations across 10 counties (Supplementary Table 2). It was predominant in 16 populations across seven regions, exceeding 95% dominance in sheep-derived populations in Southern Scotland and the West of Scotland (Supplementary Table 1).

**Fig. 1.**
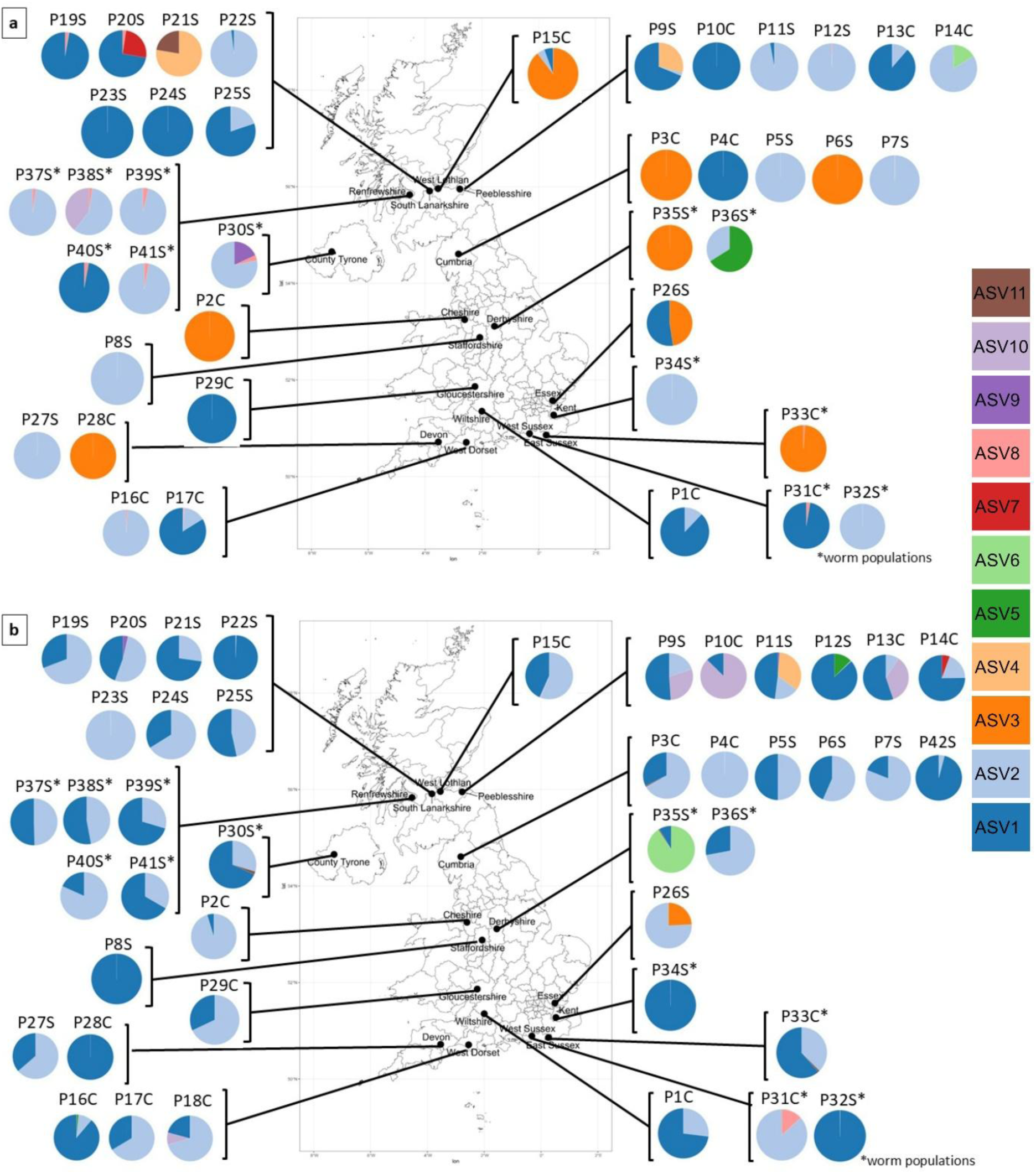
The relative frequencies of mt-ND1 and mt-COX1 ASVs in *F. hepatica* from sheep (S) and cattle (C) across 17 counties in the UK. (a) mt-ND1 ASVs and (b) mt-COX1 ASVs in 40 and 42 *F. hepatica* populations, respectively. Each pie chart represents a distinct population, originating either from adult worms (indicated with an asterisk *) or eggs purified from faeces. Individual ASVs are distinguished by different colours within the charts. The size and composition of each pie chart reflect the proportional distribution of ASVs within the respective population. Furthermore, each population is mapped to its geographical collection site, providing a visual representation of ASV diversity and distribution across the UK.

ASV2 was the second most abundant variant (35.9% of total reads), found in 24 populations across 13 counties (Supplementary Table 2) covering nine regions. It was predominant in 15 populations, reaching greater than 99% in West Midlands England and South-East England sheep flocks (Supplementary Table 1). ASV3 ranked third (14.7% of total reads), occurring in 9 populations in eight counties (Supplementary Table 2) and in seven populations, being the only ASV present in cattle and sheep fluke in North West England, as well as in sheep fluke in South West England and the East Midlands (Supplementary Table 1).

In South-East and North West England, ASV3 and ASV1 predominated in fluke populations in cattle and in South East England in fluke populations in sheep ASV2 predominated, with minor contributions from ASV8. East of England stands out for about equal proportions of ASV1 and ASV3 suggesting within-farm genetic mixing. (Supplementary Table 1).

In the Scottish Borders, ASV1 and ASV2 predominated in cattle fluke with ASV6 a rare variant and sheep fluke populations had ASV1 (68.66%) and ASV4 (28.98%) as the major contributors, and ASV2 and ASV8 as minor contributors. Southeastern Scotland cattle fluke exhibited ASV3 dominance (89.19%), with minor proportions of ASV1 and ASV2. In the West of Scotland, most sheep fluke populations ASV2 was most common and one population had ASV10 (39.52%), a variant otherwise absent from the other populations. In Northern Ireland’s County Tyrone, fluke in a single sheep population predominantly showed ASV2 (78.16%), and other varients were ASV9 (18.15%) and ASV8 (3.69%), (Fig. 1a, Supplementary Table 1).

Notably, some ASVs were found in some specific regions and fluke populations. For example, ASV4 was found locally in two Southern Scotland sheep populations (P21S, 77.62% (dominant); P25S, <0.1% (rare)). Similarly, ASV6 was rare but reached 16.1% in a single Scottish Borders cattle population (P14C). ASV7 reached a high frequency (25.41%) in a single Southern Scotland sheep flock (P20S). ASV8 was a widely distributed but rare variant observed at ≤3% abundance but present across multiple regions. Moreover, ASV9 was primarily found in Northern Ireland, while ASV10 was detected in the West of Scotland sheep flock, indicating geographically isolated variants (Supplementary Table 1).

### Network trees clustering analysis of the mt-ND-1 locus

The Neighbour-Net network tree with pie charts confirmed a highly connected parasite populations dominated by two central, multi-regional ASVs, including ASV1 (the largest node) and ASV2 (the second-largest node), which were present in both cattle and sheep fluke across multiple regions (Fig. 2a and b). ASV1 and ASV2 were connected via ASV3 or low-abundance ASV8, a genetic link between England and Scotland cattle fluke-derived dominant ASVs and Scotland sheep fluke-derived dominant ASVs. Peripheral ASVs for ASV1 included ASV5, ASV6, ASV9, ASV10, and ASV11. Peripheral ASVs for ASV2 were ASV4 and ASV7. Notably, all 11 ASVs were found in sheep fluke. In comparison, four ASVs (ASV10, ASV11, ASV5, and ASV7) were not detected in cattle fluke (Fig. 2b). Median Joining Network tree of mt-ND1 (Supplementary Fig. 3a) showed similar linkages among ASVs to the Neighbour-Net tree (Fig. 2).

**Fig. 2.**
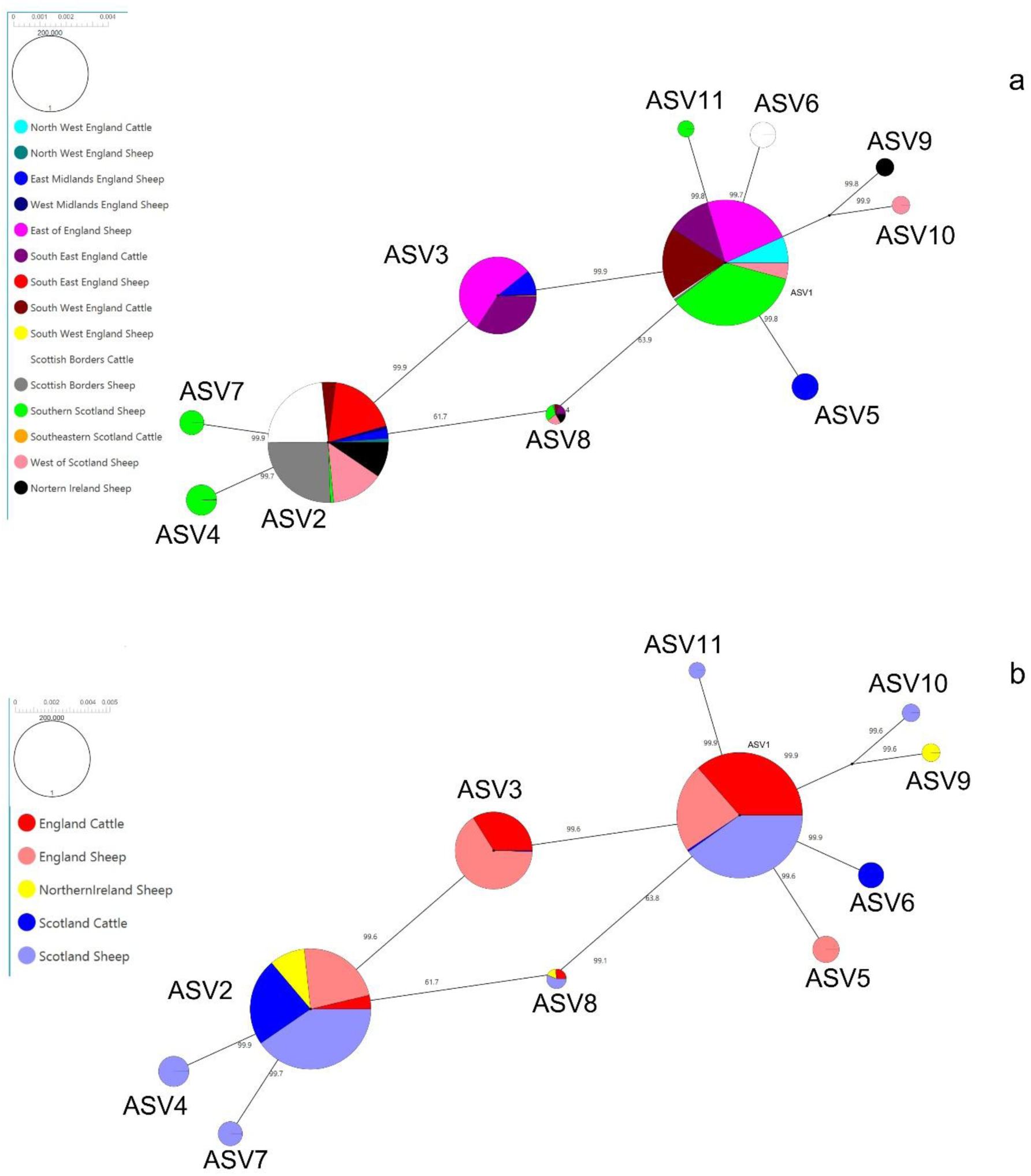
Network tree and clustering of all ASVs based on region and host (a) mt**-**ND1 Neighbour Net tree in Split tree. Each pie chart shows regions represented by different colours, with representative ASVs displayed. The pie chart represents the ASV distribution, and its frequency in all populations found in the region. The branch lengths were calculated using the HKY85 Distance method, as determined to be best by jModeltest 2.1.10. (b) Each pie chart presents the countries of the UK, represented by different colours: England cattle (red), England sheep (light red), Northern Ireland sheep (yellow), and Scotland cattle (blue), and Scotland sheep (light blue), where representative ASVs were recorded. The pie chart shows the ASV distribution and its read frequency across all populations in the countries.

The mt-ND1 PCA plot based on sequence reads of 11 ASVs from 40 populations showed partial clustering of *F. hepatica* populations by host and region, with the first two principal components explaining PC1 (18.31%) and PC2 (15.18%) of the total sequence read data (33.34%) (Fig. 3a). Populations of *F. hepatica* in sheep and cattle across the UK showed both substantial overlap and some degree of regional clustering, indicating high gene flow with occasional location-specific patterns. Most sheep derived populations fell into Cluster 1, from 9 geographical regions including South East England, East Midlands England, South West England, North West England, West Midlands England, Scottish Borders, Southern Scotland, West of Scotland, and Northern Ireland. There were two regions not found in cluster 1 including East of England, and Southeastern Scotland. This suggests that sheep across different areas carry genetically similar parasite variants. Sheep fluke populations are more evenly distributed between Cluster 1 and Cluster 2. In Cluster 2, the parasite population in sheep and cattle was widely distributed in 9 regions however, samples from West Midlands England and Northern Ireland were not represented in this cluster. This indicates that the parasite variants are not geographically restricted and are prevalent in both host species.

**Fig. 3.**
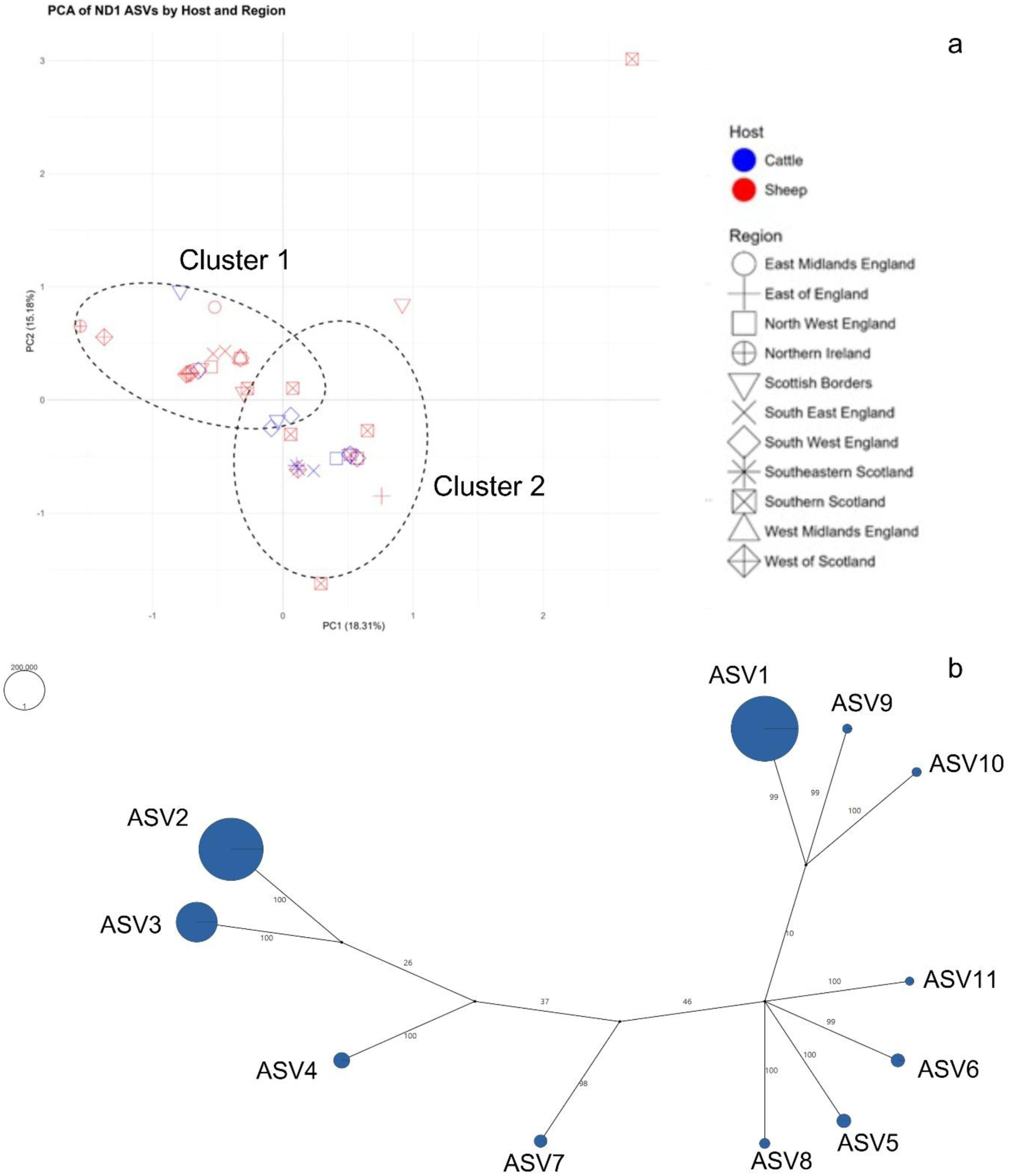
Principal Component Analysis (PCA) plot and topology tree of all ASVs based on region and host. (a) PCA of *F. hepatica* populations based on 11 mt-ND1 ASV sequence abundance. Each point represents a population, with symbols representing the different regions and colour the host species. The axes represent the first two principal components (PC1 and PC2), which explain 18.31% and 15.18% of the variance, respectively. (b) Split topology tree of mt-ND1 with the UPGMA method. The pie chart circles in the tree represent the frequency of ASVs sequence reads in all populations.

A few populations demonstrated regional genetic divergence. For example, one sheep fluke population from Southern Scotland (P21S) clustered apart with very high PC1 and PC2 values, suggesting unique ASVs in the area. Similarly, Scottish Borders sheep (P9S) and Southern Scotland sheep (P20S) also showed genetic separation from the main clusters.

The split topology tree of mt-ND1 sequences showed that one cluster is dominated by ASV1 and groups closely with ASVs 9 and 10. The second cluster is defined by ASV2 and ASV3, indicating an evolutionary relationship between these variants. Remaining ASVs (ASV4–ASV11) are distributed along shorter branches in between main variants, representing low-frequency variants that are genetically closer to one of the two dominant clusters (Fig. 3b).

### Genetic diversity analysis of the mt-ND-1 locus

The analysis of molecular variance (AMOVA) from PopArt revealed that the majority of genetic variation in *F. hepatica* mt-ND1 populations occurred within populations (137.85%). Variation among groups accounted for only 1.31% of the total, and variation among populations was negative (–39.17%) (Table 1, Supplementary Fig. 3a). The negative variance detected in the AMOVA results showed lack of genetic differentiation and should be interpreted as zero [37,38]. All fixation indices were low and non-significant (Phi ST = – 0.3785, *P* = 1.000; Phi SC = –0.3969, *P* = 0.997; Phi CT = 0.0131, *P* = 0.544), confirming the absence of significant genetic differentiation between regions or populations. Similar AMOVA results were confirmed by Arlequin (Supplementary File 1). These values suggest high genetic connectivity and gene flow among populations across the UK, consistent with the sharing of common ASVs between regions and hosts. Overall, nucleotide diversity (π=0.00502) indicated low genetic variation in the population. Tajima’s D neutrality test was not significant -0.372, (*p* = 0.623).

**Table 1.**
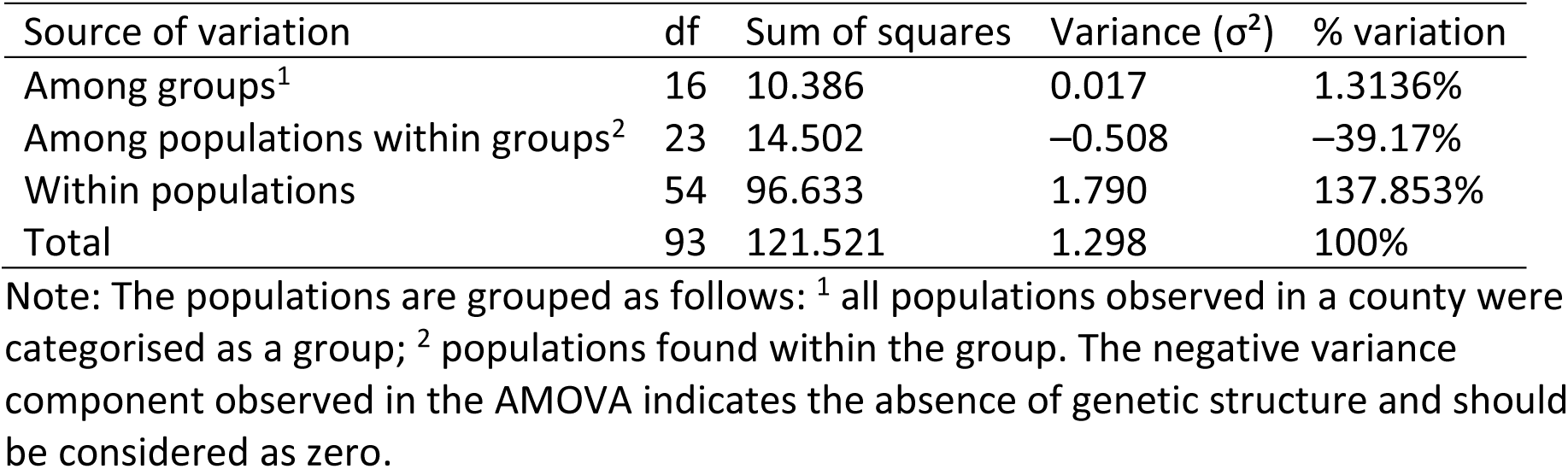
AMOVA for 11 ASVs of mt-ND1.

### Geographical distribution of mt-COX1 locus ASVs of *F. hepatica*

A total of 11 ASVs (312 bp to 319 bp) (accession numbers PX861700-PX861710) were identified at the *F. hepatica* mt-COX1 locus, using reference sequences of mt-COX1 (Supplementary Fig. 4), and their frequencies were recorded in 42 populations in different counties of the UK (Fig. 1b, and Supplementary Table 3). A total of 1.764 million sequence reads were generated across the 42 parasite populations, of which 1.33 million reads (75.2%) were from two predominant ASVs and 437,229 (24.8%) reads corresponded to rare ASVs (Supplementary Table 3). ASV1 and ASV2 were the dominant variants circulating in cattle and sheep fluke in different regions. The rare ASV3 to ASV11 were often geographically restricted.

The most abundant variant was ASV1, contributing 45.0% of sequence reads overall and detected in 41 populations across all 17 counties (Supplementary Table 4). It was the predominant variant found both in cattle and sheep fluke in 22 populations, frequently representing 75% of sequence reads. For example, in sheep fluke, ASV1 was observed in the West Midlands England (100% of reads), South East England (>99%), North West England (96%), Southern Scotland (99.46%), in the West of Scotland (>70%), Scottish Borders (86.87%) and a major variant in Northern Ireland (68.23%). In cattle, ASV1 was common in South West England (88.81%) and the Scottish Borders (75.2%). In North West England, ASV1 and ASV2 were found in equal proportions of 50% each (Supplementary Table 3). ASV2 ranked second in abundance, with overall sequence reads of 45.1% in 38 populations and 16 counties. ASV2 was found to be dominant in cattle and sheep fluke across 18 populations from 8 regions. It was widespread in cattle fluke from the North West (>95%), South West (>66.34%), and South East England (86.67%). ASV2 was highly prevalent in sheep fluke across regions of the UK including North West England (81.07%), East of England (75.77%), the West of Scotland (81.69%) and Southern Scotland (>52.06%) (Supplementary Table 3). Of the other variants only ASV6 and ASV10 were common. ASV6 predominated in East Midlands sheep fluke (90.95%), and ASV10 was predominant in Scottish Borders cattle fluke (87.5%). The remaining ASVs were present only as rare variants in different populations (Supplementary Table 3).

Although ASV1 and ASV2 were found to be predominant, the analysis of rare variants highlighted their presence in many regions, including Northwest England, Southwest England, Southern Scotland, and South Lanarkshire, ranging from 0.04% to 46.5% of sequence reads. There were certain rare variants noted in sheep and cattle fluke, for example, in sheep fluke ASV3 (24.2%) was in the East of England, ASV4 (34.01%) and ASV5 (12.64%) appeared in Scottish Borders, and ASV7 (<0.01%) in East of England. In cattle, ASV3 (0.04%) in South East England ASV5 (1.31%) in South West England, ASV7 (5.51%) occurred in Scottish Borders, and ASV8 was noted 13.29% and <0.01% in South East England and Scottish Borders, respectively. There were rare variants found in both hosts including ASV9 (< 4%) in South West England, East Midlands England, Scottish Borders, South Lanarkshire, and West of Scotland. ASV10 (0.53% to 29.3%) appeared in Scottish Borders, South Eastern Scotland, and in South West England, while ASV11 (<0.01% to 2.08%) in North West England, Southern Scotland, Northern Ireland, South East England, and East Midlands (Supplementary Table 3). Overall, the rare ASVs demonstrated regional and host-specific patterns and may emerge in the future in both sheep and cattle populations across the UK.

### Network trees clustering analysis of the mt-COX1 locus

The Neighbour Net analysis of the 11 mt-COX1 ASVs revealed two main nodes. ASV1 and ASV2 were connected through the low-abundance ASVs 4, 6, 10, and 11, forming a genetic bridge between host- and region-specific ASVs (Fig. 4a). The peripheral ASVs for ASV1 included ASV3, ASV7, ASV8, and ASV9. For ASV2, there was only ASV5. For instance, ASV1 was most abundant in sheep fluke across Scotland and England, but was also found in considerable amounts in cattle across the same regions. ASV2 was detected in sheep and cattle fluke in England, as well as in sheep fluke in Scotland, with low abundance in sheep fluke from Northern Ireland and in cattle fluke from Scotland. In contrast, other ASVs showed strong geographic and host specificity. ASV3 and ASV6 were found in English sheep, ASV4, ASV5, and ASV9 were found in Scottish sheep, ASV7 was specific to Scottish cattle, ASV8 was restricted to English cattle, and ASV11 was found mainly in Northern Ireland sheep (Fig. 4b).

**Fig. 4.**
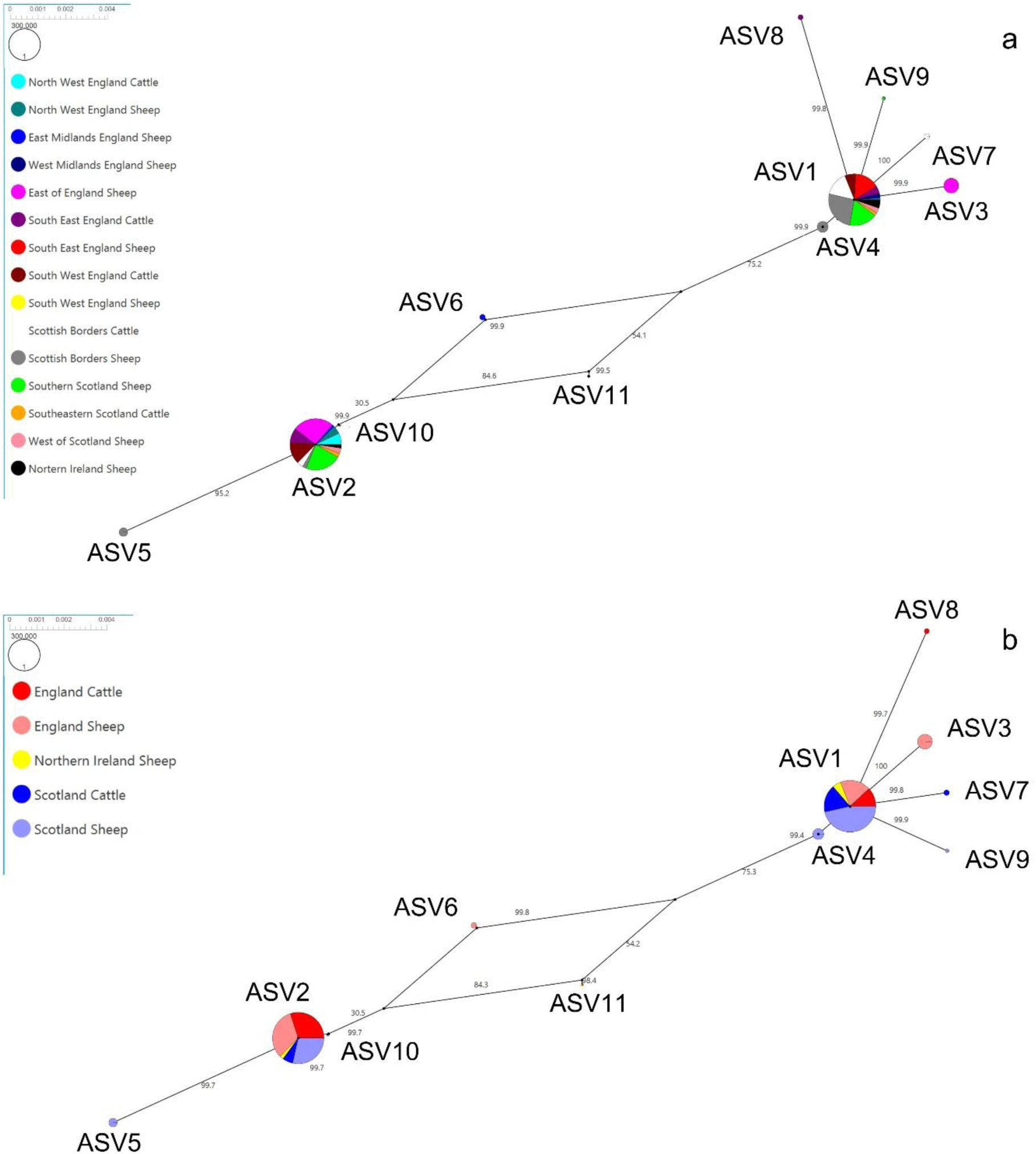
Network tree and clustering of all ASVs based on region and host (a) mt-COX1 Neighbor Net tree in Split tree. Each pie chart shows regions represented by different colours, with representative ASVs recorded. The pie chart circle represents the ASV distribution, and its sequence reads frequency in all populations found in the region. The branch lengths were calculated using the HKY85 Distance method, as determined to be best by jModeltest 2.1.10. (b) Each pie chart shows the countries of the UK, represented by different colours: England cattle (red), England sheep (light red), Northern Ireland sheep (yellow), and Scotland cattle (blue) and Scotland sheep (light blue), with representative ASVs recorded. The pie chart shows the ASV distribution and read frequency across all populations in the countries.

Median Joining Network tree of mt-COX1 (Supplementary Fig. 3b) showed similar linkages among ASVs as the Neighbour Net tree (Fig. 4a and b).

The PCA plot based on mt-COX1 sequence reads from 42 *F. hepatica* populations showed a single cluster by host and region, with the first two principal components explaining PC1 19.07% (PC1) and 15.69%(PC2) of the total (34.76%) (Fig. 5a). All parasite populations in sheep and cattle fluke across the UK show substantial overlap and indicating high gene flow. The topology tree of 11 mt-COX1 ASVs showed two major ASVs (ASV1 and ASV2) at the ends of the tree, and the remaining ASVs were mainly distributed along shorter branches in between the main ASVs, indicating phylogenetic separation among ASVs (Fig. 5b).

**Fig. 5.**
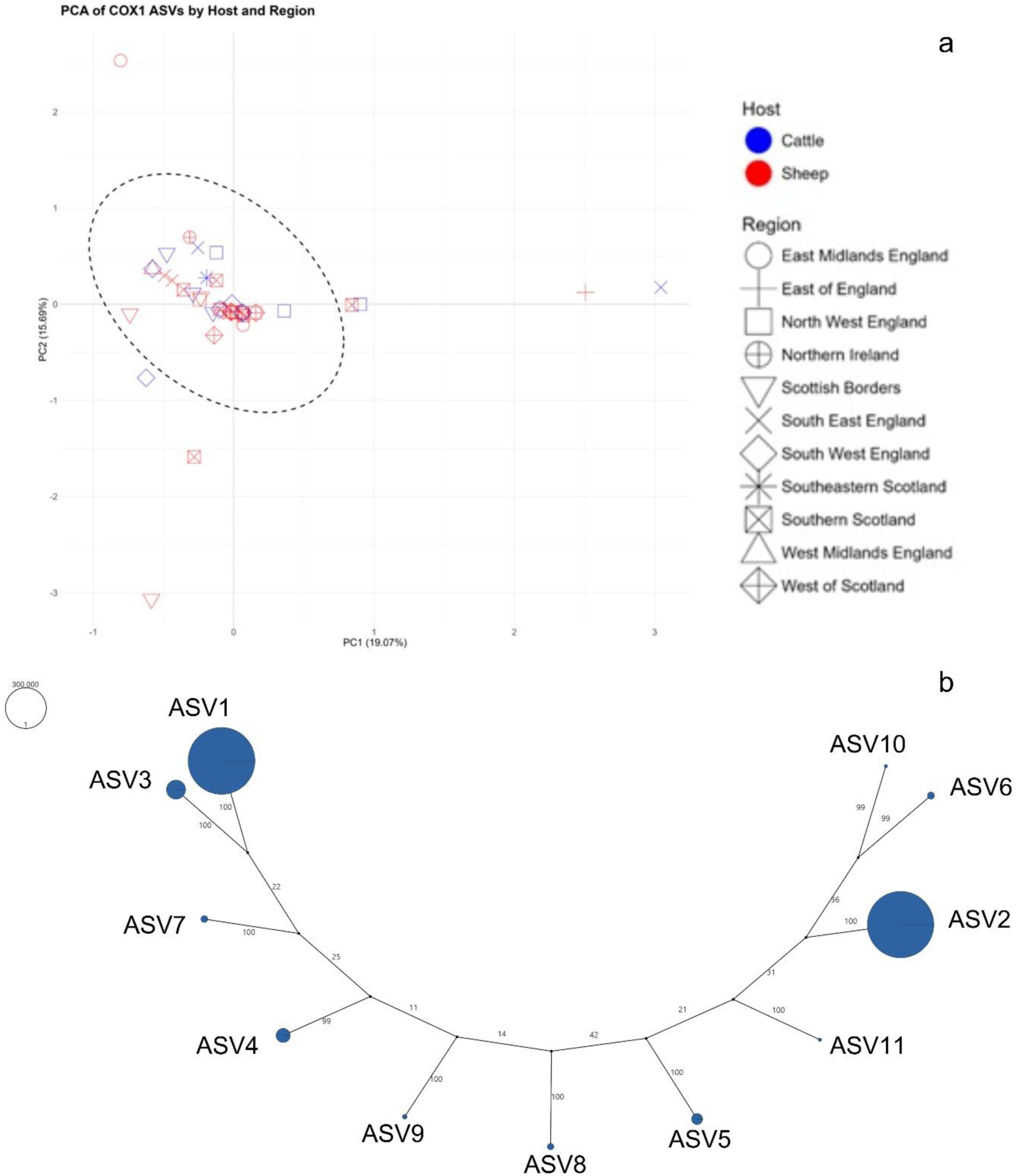
Principal Component Analysis (PCA) plot and topology tree of all mt-COX1 ASVs (a) PCA of *F. hepatica* populations based on 11 mt-COX1 ASV sequence abundance. Each point represents a population, with symbols representing the different regions and colour the host species. The axes represent the first two principal components (PC1 and PC2), which explain 19.07% and 15.69% of the variance, respectively. (b) Split topology tree of mt-COX1 using the UPGMA method. The pie charts represent the frequency of ASVs’ sequence reads in all populations.

### Genetic diversity analysis of the mt-COX1 locus

The AMOVA results showed that the genetic variation occurred within populations (139.4%), while variation among groups (0.75%) and among populations within groups (−40.2%) was very low (Table 2, Supplementary Fig. 3b). Similar AMOVA results were found using Arlequin (Supplementary File 2). Consistently, fixation indices were very low or negative (Phi ST = −0.394, Phi SC = −0.405, Phi CT = 0.007), and none were statistically significant (*p* > 0.05). Together, these results suggest that there is no significant genetic variation among groups and populations. Genetic diversity was found only within populations, not across hosts or geographic regions. Overall, nucleotide diversity (π = 0.00834) was low across all populations. Tajima’s D test was 0.375 and not significant (*p* = 0.343).

**Table 2.** AMOVA for 11 AVS of mt-COX1.

| Source of Variation | df | Sum of Squares | Variance ($\sigma^2$) | % Variation |
| --- | --- | --- | --- | --- |
| Among groups<sup>1</sup> | 16 | 29.009 | 0.040 | 0.74717 % |
| Among populations within groups<sup>2</sup> | 25 | 50.794 | -2.153 | -40.16418 % |
| Within populations | 68 | 508.133 | 7.473 | 139.41701 % |
| Total | 109 | 587.936 | 5.360 | 100 % |

Note: The populations are grouped as follows: <sup>1</sup> all populations observed in a county were categorised as a group; <sup>2</sup> populations found within the group. The negative variance component observed in the AMOVA indicates the absence of genetic structure and should be considered as zero.

## Discussion

The present study developed a metabarcoding approach using multiplex mitochondrial markers targeting the mt-ND1 and mt-COX1 loci to study the population genetics of *F. hepatica*, using samples from natural infections collected across different areas of the UK. The application of the metabarcoded multiplexed markers in deep amplicon sequencing can provide an efficient tool for investigating genetic diversity patterns in multiple populations of *F. hepatica* that can further inform transmission dynamics and infection rates. This study demonstrated that *F. hepatica* populations with a small number of predominant ASVs were circulating in both cattle and sheep with high gene flow across different regions of the UK. Alongside this a few rare geographically restricted ASVs were also noted. The results highlighted a pattern of widespread ASV flow across the UK, which may be driven by clonal propagation of the parasite in intermediate snail hosts, livestock movement, grazing practices, and parasite adaptation to the UK environment.

This study utilised mitochondrial markers to analyse the genetic diversity of *F. hepatica* populations because mitochondrial DNA is useful for population genetics investigations, due to its high copy number [33], maternal inheritance [32], and transmission without recombination under neutral heteroplasmy conditions [35]. These properties make mitochondrial DNA a powerful target for investigating evolutionary studies. Mitochondrial markers have been used to study genetic diversity and infection dynamics in *F. gigantica,* targeting the mt-ND1 locus where a single mt-ND1 variant of *F. gigantica* was predominant in most of the hosts in Pakistan [12], and to investigate *C. daubneyi*, using the mt-COX1 locus. Notably, multiple variants of *C. daubneyi* infections, were detected across different geographic regions of the UK [17]. A study from Malawi supported the suitability of both mt-ND1 and mt-COX1 loci for population genetics studies in *F. gigantica* and reported recent population expansion [22]. Other studies also supported the use of mitochondrial markers for genetic diversity and gene flow studies in *F. hepatica* [39–42].

The amplification success across 90 samples, with 78 and 84 yielding sequence reads for mt-ND1 and mt-COX1, respectively, highlighted the effectiveness of our method. Further, the deep amplicon sequencing technique detected both dominant and rare ASVs, enhancing our understanding of parasite population genetics. This ability to recover ASVs in different parasite populations across geographical regions of the UK demonstrated that multiplex PCR combined with next-generation sequencing can be a valuable tool for studying fine-scale genetic structure. Previous studies used PCR-RFLP, conventional PCR, and Sanger sequencing to amplify and analyse the mitochondrial genome regions to investigate genetic variations [22,43,44]. However, these methods are low-throughput, time-consuming, relatively expensive when handling medium to high numbers of samples [45]. In contrast, high-throughput deep amplicon sequencing using the Illumina MiSeq platform offers a convenient and cost-effective method [46] for handling a medium to high number of samples, with thousands and millions of sequence reads generated per population in a single run [12,17,47].

This study identified some predominant ASVs across different regions of the UK, as well as more locally restricted variants. For instance, the most common three variants of mt-ND1 (ASV1-3) were widely distributed across multiple regions and hosts and together accounted for over 80% of infections. Similarly, two mt-COX1 variants were dominant. AMOVA results showed that most genetic variation occurred within populations rather than between counties or regions, which is consistent with high level of gene flow. Based on microsatellite genotyping, previous research in the UK also found high genetic diversity and gene flow, as well as an absence of defined population structures. This was believed to be due to the clonal emergence of *F. hepatica* infections through the intermediate snail host [13], such that a single miracidium infecting a G. *truncatula* mud snail can generate multiple genetically identical cercariae [13]. However, metacercariae may mix on pasture, resulting in a more varied genetic profile before ingestion by the definitive host [12].

Findings from *F. hepatica* isolates from three geographical regions of China supported our results, showing that genetic variation can occur within populations rather than between populations [48]. In another example from the European context, in the Netherlands, genetic diversity was also reported mostly within populations rather than between populations [49]. In Algeria, low genetic diversity and a common origin for the parasite’s countrywide distribution were reported, with only two variants from the mt-COX1 gene [50]. In Colombia, no genetic diversity was found among *F. hepatica* parasites. However, the authors mentioned that this might be due to the low resolution of the molecular markers used including nuclear markers (28S, β-tubulin 3, ITS1, ITS2), and mitochondrial marker (mt-COX1) [51].

In contrast, high genetic diversity between populations was found in German dairy cattle using mitochondrial (mt-ND1 and mt-COX1) and eight microsatellite markers [44], in cattle and sheep in Spain and Peru with mt-ND1 marker [52], and in grazing cattle in Australia using mt-ND1 and mt-COX1 markers [42]. Further, a high number of mt-COX1 mitochondrial variants have been reported in cattle and horses in Chile [53]. A 2007 study reported that a single animal in Ireland could harbour ten distinct mitochondrial variants of *F. hepatica*, and the author related that this genetic diversity predates the last ice age [43]. A global-scale analysis of NCBI data showed that both mt-ND1 and mt-COX1 locus-specific variants were circulating in different parts of the world, with high Tajima’s D values and a low likelihood of future population growth [54]. However, from Armenia, Algeria, Brazil, Spain, and Ecuador, negatively significant Tajima’s D values were reported for mt-ND1 along with mt-COX1 showed deviation from neutrality, supporting recent population expansion [54]. This contrast between the neutral, locally mixed parasite populations found in this work from UK and the variable patterns observed globally highlights how local gene flow can influence in future evolutionary processes in *F. hepatica* populations.

Unlike the predominant ASVs, the rare ASVs identified in this study exhibited a mostly regional distribution, with only a few rare ASVs found in multiple populations across different areas, for example, mt-ND1 (ASV8) and mt-COX1 (ASV9-ASV11). This showed the emergence of localised variants, potentially linked to specific environmental factors, especially environmental temperature and soil conditions, which can influence the transmission dynamics of *F. hepatica* [55]. For example, *G. truncatula* egg masses are rarely observed in shaded areas [56], and this could affect opportunities for fluke transmission in different environments. *Fasciola* eDNA was less frequently detected in dark brown or black soils, which are rich in organic matter in the form of peat and have lower pH values [56]. This could impact the distribution of intermediate snail hosts and the clonal expansion of rarer ASVs. Further, *Fasciola* spp. egg hatching and development are optimal between 20 °C and 30 °C and inhibited at temperatures below 10 °C. The time required for miracidia hatching decreased with increasing temperature, and shedding of cercariae from snail hosts was most rapid at around 27 °C [55]. Viability of metacercariae declined at higher temperatures but could be prolonged under high humidity. Snails grow best at 25 °C, and their susceptibility to *Fasciola* infection is also temperature dependent [55]. Climate variation and change could therefore impact the geographic distribution of liver fluke, and drive adaptation to local conditions. For instance, *F. hepatica* infections have historically been low in regions such as Southern Europe, but climate shifts may increase winter risk due to temperature and moisture fluctuations falling into suitable ranges [11]. Increased fluke infections have been recorded in the EU and the Northern Altiplano in South America [57]. Additionally, a study in New Zealand utilised historic climate data (1972-2012) and predicted that areas with low initial risk, such as Canterbury and Otago, could see a near 200% rise by 2090 [58]. Although we have currently identified rare and locally restricted ASVs, these studies highlight how they could serve as potential reservoirs of future genetic diversity that could become epidemiologically significant under shifting environmental conditions. Further research is needed to determine how genetic diversity measured using different markers to differences in biological responses and environmental conditions by *F. hepatica* and other parasites.

In our study, Neighbour-Net, median joining networks and PCA showed that UK *F. hepatica* populations are interconnected and dominated by a few ASVs. In the PCA plot, mt-ND1 revealed one cluster containing mainly sheep-hosted parasite populations and a second cluster containing both sheep and cattle-hosted populations. In contrast, for mt-COX1, parasite populations from both sheep and cattle across different regions of the UK formed a single, overlapping cluster. Our study did not gather data on co-grazing of sheep and cattle. However, sheep and cattle often co-graze in Northern Ireland and may be infected with the clonal variants of parasites [43]. Prichard et al., 2005 also attributed fluke outbreaks in East England to co-grazing with sheep imported from other areas and high rainfall during summer [59]. This homogenisation of ASVs across populations in England, Scotland, and Northern Ireland is likely due to livestock movement, a common practice in the UK agricultural system and important for the spread of various diseases [60,61]. The role of animal movement in parasite transmission and high gene flow has been well documented [12,13,17,62]. Furthermore, greater genetic structure was observed in parasite populations in sheep rather than cattle, which may be due to differences in grazing behaviour. Sheep tend to feed closer to soil and waterlogged areas than cattle, potentially increasing exposure [63]. In contrast, higher genetic diversity was reported in cattle fluke than in sheep and goat fluke in Iran [64], but this may be because the prevalence of this disease is higher in cattle than in sheep in Iran [65,66]

The present study has limitations. The use of only two mitochondrial markers (mt-ND1 and mt-COX1) provided valuable resolution but captures only maternally inherited variation. The use of nuclear markers or whole-genome data could further enhance understanding of nuclear-level population genetics, and host adaptation [13,23,67]. Although 90 samples from 17 counties were analysed, the sample coverage was uneven, with some regions represented by only one or two populations, and no faecal samples were obtained from Northern Ireland. Unequal sampling can overrepresent genetic structuring and overstate the apparent dominance of a few ASVs in different regions. Finally, intermediate host snail and livestock movement data were not included in this study, although both are known to influence the spread of *F. hepatica* infection strongly.

Future research should aim to overcome the stated limitations by optimising nuclear markers and whole-genome sequencing to complement mitochondrial markers and capture genetic polymorphisms and adaptive behaviours (manuscript in preparation). A higher number of samples across multiple years and seasons would help to track deeper dynamics of ASV diversity. Linking parasite genetic data with phenotypic outcomes, such as flukicide efficacy through faecal egg count reduction tests, infection intensity, and productivity losses in cattle and sheep, will strengthen knowledge of fluke epidemiology and control options. Assessment of genetic diversity of *F. hepatica* in intermediate snail hosts will provide insights into bottlenecks and parasite persistence in wetlands, while combining parasite genetics with livestock trade, and grazing movement data will enable causal inference about how gene flow is maintained across the UK. These approaches can generate more understanding of *F. hepatica* transmission and evolution, supporting targeted interventions against fasciolosis. The methods optimised and described here provide an additional tool for collecting genetic data and linking it with infection and disease outcomes, as well as inferring patterns of parasite transmission and spread.

## Conclusion

In conclusion, a multiplex mitochondrial metabarcoding approach has been developed here, providing a platform for medium to large-scale population genetic studies of *F. hepatica* infection. This study demonstrated that *F. hepatica* populations in the UK are largely genetically interconnected and dominated by a small number of widespread variants in both cattle and sheep. The findings confirm that cross-transmission of fluke between co-grazing sheep and cattle is likely, although some genotypes seem to be more restricted to sheep fluke. In addition, rare and region-specific variants were found at low frequencies, which may contribute to the future emergence of new variants in the UK if not controlled. Our findings support the idea that high gene flow can result from parasite adaptation in the UK environment alongside high levels of livestock movement. These findings are important for understanding transmission dynamics, detecting emerging variants, and informing effective control strategies for *F. hepatica* infections to livestock farmers.

## Methods

### Field samples

A total of 90 field samples comprising 78 faecal egg samples and 12 adult worm samples were selected, which identified as *F. hepatica* positive in our previous study [68]. Multiple samples collected from the same sheep and cattle farms, veterinary practitioners, or counties within close timeframes were merged into single parasite populations. Additionally, adult fluke obtained from abattoirs and at post-mortem examination from the same animal were treated as a single population. All populations were assigned by host species, cattle (n=15) and sheep (n=27) for downstream analyses.

The samples were collected across 17 counties in the UK between December 2022 and May 2024 in collaboration with cattle and sheep farmers, as well as registered veterinary practitioners, in accordance with ethical approval NASPA-2122-04 [68]. The populations were further categorised into 11 regions across the UK including North West England (Cheshire, Cumbria), East Midlands England (Derbyshire), West Midlands England (Staffordshire), East of England (Essex), South East England (East Sussex, Kent, West Sussex), South West England (Dorset, Devon, Gloucestershire, Wiltshire), Scottish Borders (Peeblesshire), Southern Scotland (South Lanarkshire), Southeastern Scotland (West Lothian), West of Scotland (Renfrewshire), Northern Ireland (County Tyrone). Adult worm populations were obtained from the livers of infected animals at abattoirs described [68]. All samples were transported to the School of Veterinary Medicine at the University of Surrey and stored at –20°C for subsequent analysis.

### DNA extraction

DNA was extracted from a pooled sample of head tissue obtained from all available adult worms per host and from faecal egg samples, following the methods described [68]. Elution was performed in 50 μL of molecular biology-grade water (Cytiva HyClone™), and the eluate was stored at –80 °C for subsequent analyses.

### Development of multiplex mitochondrial markers

PCR was conducted using *Fasciola* genus-specific mt-ND1 primers, resulting in a product size of 311 bp [12]. In addition, mt-COX1 primers for *F. hepatica* were designed and tested in this work (Supplementary Table 5), potentially aiming for a product length of 319-475 bp. These primers were designed by aligning 435 mt-COX1 sequences from *F. hepatica* available on Genbank and selecting a region showing variations and a conserved region after visualising different sequences using *Primer3* in Geneious Prime version 8.0.5. The primer pair was 20 bp lengths, Tm values range was 59.2-59.8 °C, GC content 50–60%, and a forward and reverse Tm difference of 0.6 °C. Primers did not contain any hairpins, self-dimers, and cross-dimers.

PCR conditions were first optimised using positive control DNA in a gradient PCR with annealing temperatures ranging from 52°C to 60°C for amplification of mt-COX1 (Supplementary Fig. 1c). PCR was performed in duplicate and twice with 2 μl of the DNA template using DreamTaq Green PCR master mix (Thermo Scientific, USA) in a 25 μl reaction mix and primer concentrations of 200 nM in the final volume. Final PCR conditions included 35 cycles of initial denaturation at 95°C for 5 minutes, followed by denaturation at 95°C for 1 minute, annealing of mt-ND1 primers at 50°C for 1 minute and mt-COX1 primers at 55°C for 1 minute, and extension at 72°C for 1 minute, with a final extension at 72°C for 5 minutes. Water was used as a no-template control. The resulting PCR products were processed for Sanger sequencing.

A multiplex PCR was developed using both mt-ND1 and the mt-COX1 primers, with DreamTaq Green PCR master mix (Thermo Scientific, USA) with primer concentrations of 200 nM in 25 μl final reaction volume. The PCR was conducted for 35 cycles with conditions as follows: an initial denaturation step at 95°C for 5 minutes, followed by denaturation at 95°C for 1 minute, annealing at 53°C for 1 minute, extension at 72°C for 1 minute and a final extension at 72°C for 5 minutes. The multiplex PCR reaction was carried out in duplicate using two μl of positive control *F. hepatica* DNA, and water as a negative control.

### Deep amplicon sequencing of multiplexed mt-ND1 and mt-COX1

The metabarcoded mt-ND1 [12] and mt-COX1 (Supplementary Table 1) markers were used to target the mitochondrial DNA of *F. hepatica*. A multiplexed first-round PCR was carried out using the KAPA HiFi PCR Kit (KAPA Biosystems, South Africa) as described [12]. The second-round primer sets, adaptors, barcoded PCR amplifications, magnetic bead purification, and final library quantification were based on previously described methods [12,17,69].

### Demultiplexing of mt-ND1 and mt-COX1 sequences and bioinformatics analysis

The Illumina MiSeq system demultiplexed the sequencing data based on sample-specific barcoded indices (Supplementary Table 6), generating corresponding FASTQ files for each sample (NCBI Bioproject: PRJNA1402908, accession No: SAMN54606237-SAMN54606325, https://data.mendeley.com/datasets/822rxwph9t/1). The resulting FASTQ files were further analysed using Mothur versions 1.41.0 and 1.48.1 [70] on the University of Surrey High-Performance Computing (HPC) cluster. Sequence analysis was performed following the pipelines described in previous studies [12,17] with modifications as described below.

For reference database construction, mt-ND1 and mt-COX1 sequences were retrieved from the NCBI database for *F. hepatica* and *F. gigantica*. A total of 363 mt-ND1 sequences and 462 mt-COX1 sequences were downloaded for *F. hepatica*, which were collapsed into 79 and 97 unique reference sequences, respectively. For *F. gigantica*, 351 mt-ND1 and 337 mt-COX1 sequences were downloaded, resulting in 117 and 108 unique collapsing sequences, respectively. These reference sequence libraries for the mt-ND1 and mt-COX1 genes were used for sequence demultiplexing and alignment. The Mothur pipeline joined paired-end reads, filtered out ambiguous or low-quality sequences, and removed excessively long or short sequences. Sequences were aligned against the reference library, unique sequences were pre-clustered and abundant reads were then grouped.

ASVs were obtained from the filtered dataset using minimum read thresholds assigned to eliminate noise and sequencing artefacts using the command “split.abund”. A cutoff value of 7,000 reads per ASV was applied to the mt-ND1 dataset, and 2,000 reads per ASV were used for mt-COX1. These thresholds were determined empirically following visual inspection of the output count table files. The aligned unique sequences were split into a high- and low-abundance sequence read count table and FASTA files based on the defined cutoff value. Sequences with low abundances and below threshold were separated and discarded, while high-abundance sequences were selected for downstream analyses. Following ASV extraction for all samples, downstream processing, including sample wise sequence cleaning and sorting of FASTA files by specific ASV names, was performed in R using the phytools [71], microseq [72], biostrings [73], dplyr [74], ggh4x [75] and tidyverse [76] packages. A FASTA file and group file containing corresponding abundance data generated by Mothur were used for downstream analysis. The dataset was standardised to prevent mismatches. The data was used to generate population-specific sequence fasta files, where sequences were replicated according to their abundance values. Biostrings-based function was used to each FASTA file to improve data quality. This function removed ambiguous nucleotides (e.g., ‘N’) and non-ATCG characters and filtered out sequences shorter than 100 base pairs. The cleaned sequences were saved into new FASTA files, and ASVs were named in each sample based on the descending order of sequence read abundance. ASV sequences were visualised using Geneious version 8.0.0 (https://www.geneious.com). Finally, sample files are grouped into populations using an R script, and unique sequences were extracted for each population for downstream analysis. All R scripts and the reference sequences library used for this process are available at the Mendeley data repository (https://data.mendeley.com/datasets/822rxwph9t/1).

### Phylogenetic, network and Split Tree analysis

Phylogenetic trees were generated from unique reference sequences of mt-ND1 and mt-COX1 for *F. hepatica* and *F. gigantica*, downloaded from NCBI GenBank. The sequences were aligned using MUSCLE in Geneious v8.0.5. Further, phylogenetic trees were constructed using the Neighbour-Joining method [77]. The evolutionary distances were computed using the Maximum Composite Likelihood method [78] in MEGA11 [79] with a bootstrap value of 2000 [80].

Split trees were generated using SplitTrees4 CE 6.0.0 [81], employing the HKY85 Distance [82] and Neighbor Net method [83,84]. The most appropriate nucleotide substitution model for HKY85 Distance was identified using jModelTest 12.2.0 [85]. Split topology tree was generated using UPGMA method [86], and with the 1000 Bootstraps [80]. Moreover, the Median Joining Network tree, nucleotide diversity, Tajima D, and AMOVA analysis were performed using popart-1.7 [87,88]. AMOVA analysis was further confirmed using Arlequin [89].

## Data analysis

Pie charts, frequency distributions, and analyses of predominant and rare ASV patterns, as well as the proportional distribution of ASVs within each county and across the population, were generated to characterise *F. hepatica* populations in sheep and cattle in R using packages readxl [90], ggplot2 [91], dplyr [74], and tidyr [92]. Location data points from confirmed positive collection sites were plotted on the UK map. Geographic coordinates (longitude and latitude) for each site were taken from Google Maps (Supplementary Table 7). Mapping was performed using spatial data sourced from the UK Data Service, including Census Support Digitised Boundary Data (1840–present) and Postcode Directories (1980–present), which allowed for the accurate visualisation of ASV distribution patterns across the UK. All data analysis and visualisations were performed in R version 4.3.3 (https://cran.r-project.org/).

## Acknowledgements

Part of this work was carried out using computational HPC facilities and support provided by the Research Computing Services team within IT Services at the University of Surrey, specifically the Eureka2 HPC cluster (https://docs.pages.surrey.ac.uk/research_computing/hpc/clusters/eureka2.html).

This research was funded by the UK Research and Innovation (UKRI), Biotechnology and Biological Sciences Research Council (BBSRC) through the FoodBioSystems Doctoral Training Programme (BB/T008776/1) and by the Sir Halley Stewart Trust (3153). For Open Access, the authors have applied a Creative Commons Attribution (CC BY) public copyright license to any Author Accepted Manuscript version arising from this submission.

We sincerely acknowledge all farmers and registered veterinary practitioners in the UK and especially Dr. Iñaki Deza-Cruz (The Royal (Dick) School of Veterinary Studies and The Roslin Institute, The University of Edinburgh, Easter Bush Veterinary Centre, Midlothian, EH25 9RG) for reading the manuscript and sample collection. Dr. Sai Fingerhood (Department of Veterinary Pathology, University of Nottingham, UK), and Dr. Mark W. Robinson (School of Biological Sciences, Queen’s University Belfast, UK) for their valuable assistance in sample collection.

## Contributions

Muhammad Abbas: conceptualisation, investigation, methodology, bioinformatics, validation, visualisation, data curation and analysis, writing original draft, review and editing; Kezia Kozel: methodology, writing review and editing; Nick Selemetas: writing review and editing, supervision; Olukayode Daramola: writing review and editing, supervision; Eric R Morgan: conceptualisation, funding acquisition, supervision, writing review and editing; Umer Chaudhry: conceptualisation, writing review and editing, supervision; Martha Betson: conceptualisation, writing review and editing, supervision, funding acquisition, project administration.

## Ethical statement

Non-invasive collection of faecal samples was approved by the NASPA (Non-Animal Scientific Procedures Act) sub-committee of AWERB, University of Surrey, UK, under the reference NASPA-2122-04 for the project “Developing Novel Rapid Diagnostics for Neglected Parasitic Diseases.” Adult *F. hepatica* were collected at licensed slaughterhouses and through post-mortem examination. Completion of a University of Surrey SAGE-AR (ID 638929-638920-101535552) indicated that no formal ethical approval was required for adult fluke sampling.

## Supplementary information

Supplementary Fig. 1 to 4

Supplementary Files. 1 to 2

Supplementary Tables 1 to 7

## Rights and permissions

All sequencing data reported in the paper are available under NCBI BioProject ID PRJNA1402908 and accession numbers: SAMN54606237-SAMN54606325, PX861700-PX861710 and PX902280-PX902290.

In addition, sequence data, R script, and codes are available at the Mendeley database https://data.mendeley.com/datasets/822rxwph9t/1

All other data are reported in the paper and associated supplementary material.

## Funding

Muhammad Abbas received funding from the UK Research and Innovation (UKRI), Biotechnology and Biological Sciences Research Council (BBSRC) through the FoodBioSystems Doctoral Training Programme for project ID FBS2022 titled “New tools for sustainable control of liver fluke in ruminants” Grant Ref: BB/T008776/1. Further, this research was funded by the Sir Halley Stewart Trust under the project “Rapid Diagnostics for Neglected Parasites.

## Competing Interest

The authors declare that no financial interests or personal relationships could have influenced the work reported in this paper.

## Notes

https://data.mendeley.com/datasets/822rxwph9t/1

### Competing Interest Statement

The authors have declared no competing interest.

