## Supplementary Fig. 1 to 4 for "Limited genetic structure and high gene flow in *Fasciola hepatica* populations infecting ruminants in different geographic areas in the UK"

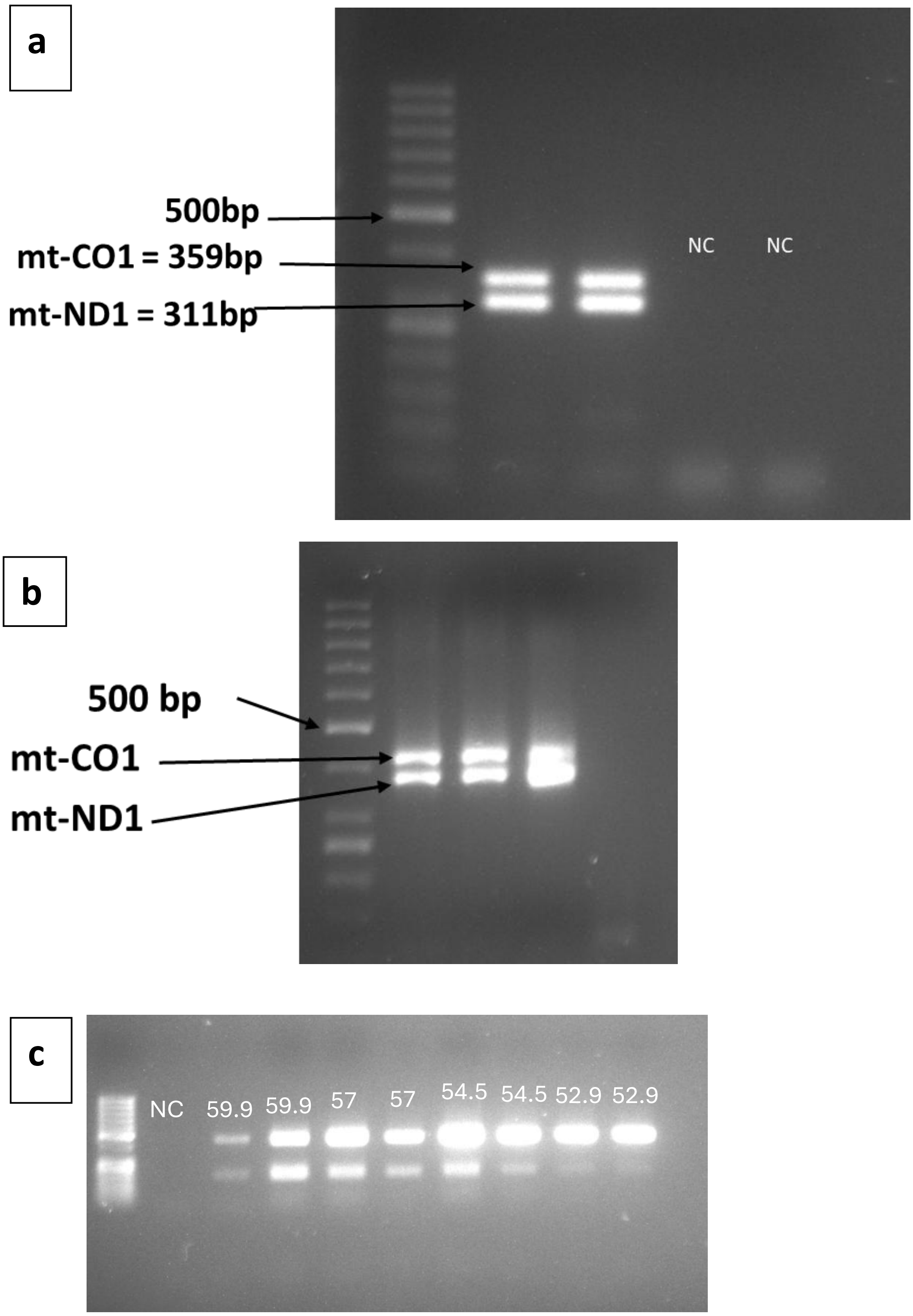


**Supplementary Fig. 1:** (a) Multiplex conventional PCR products for mt-COX1 (359bp) and mt-ND1 (311bp) compared to a 1Kb ladder (500bp marker indicated). (b) Multiplex adaptor PCR. mt-COX1 and mt-ND1 products with adaptors attached. (c) Gradient PCR using temperatures 52°C to 60°C for mt-COX1 markers to optimise annealing temperature.

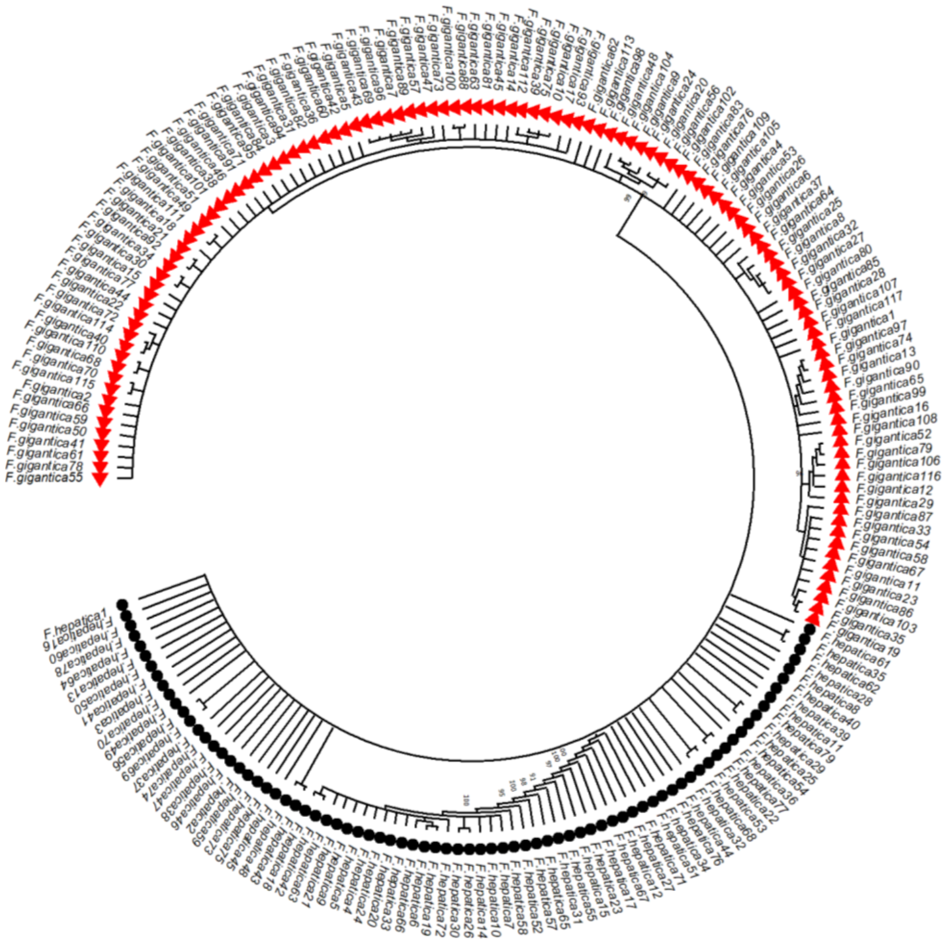

**Supplementary Fig. 2:** A Neighbour Joining tree was constructed using 2000 bootstrap replicates. *F. gigantica* and *F. hepatica* mt-ND1 loci sequences were identified through BLAST searches. In total, 196 sequences were analysed, comprising 79 *F. hepatica* sequences, represented by black circles, and 117 *F. gigantica* sequences, represented by red triangles. The two species clustered separately. Curated *F. hepatica* reference sequences were subsequently used to demultiplex and classify Illumina MiSeq mt-ND1 reads.

**a**

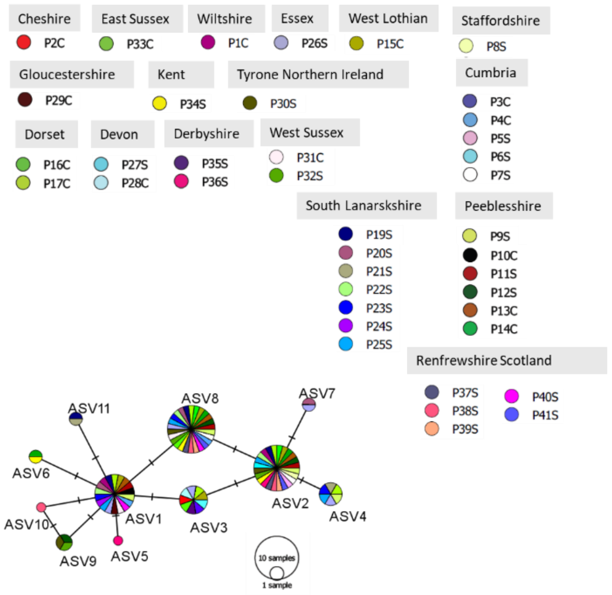


| ASV | Total Reads | Reads (%) | Population Count | Populations |
|---|---|---|---|---|
| ASV1 | 547801 | 39.0321 | 18 | P10C, P11S, P13C, P15C, P17C, P19S, P1C, P20S, P22S, P23S, P24S, P25S, P26S, P29C, P31C*, P40S*, P4C, P9S |
| ASV2 | 504737 | 35.9637 | 24 | P11S, P12S, P13C, P14C, P15C, P16C, P17C, P19S, P1C, P22S, P25S, P27S, P30S*, P32S*, P34S*, P36S*, P37S*, P38S*, P39S*, P41S*, P5S, P7S, P8S, P9S |
| ASV3 | 206597 | 14.7205 | 9 | P15C, P17C, P26S, P28C, P2C, P33C*, P35S*, P3C, P6S |
| ASV4 | 32142 | 2.2902 | 6 | P17C, P21S, P23S, P25S, P26S, P9S |
| ASV5 | 24323 | 1.7331 | 1 | P36S* |
| ASV6 | 22616 | 1.6114 | 2 | P14C, P34S* |
| ASV7 | 20472 | 1.4587 | 2 | P20S, P26S |
| ASV8 | 13509 | 0.9625 | 26 | P11S, P12S, P13C, P14C, P15C, P16C, P17C, P19S, P20S, P22S, P23S, P25S, P26S, P30S*, P31C*, P32S*, P33C*, P34S*, P37S*, P38S*, P39S*, P40S*, P41S*, P4C, P7S, P9S |
| ASV9 | 11156 | 0.7949 | 3 | P12S, P30S*, P32S* |
| ASV10 | 10913 | 0.7776 | 1 | P38S* |
| ASV11 | 9196 | 0.6552 | 2 | P19S, P21S |

**b**

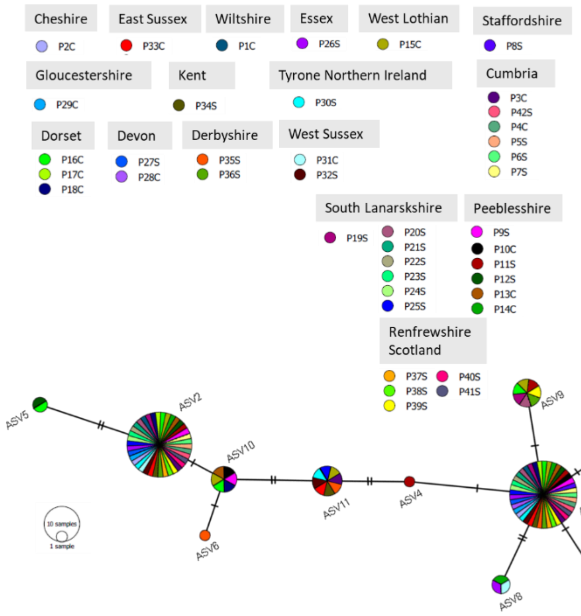


| ASV | Total Reads | Reads (%) | Population Count | Populations |
|---|---|---|---|---|
| ASV1 | 807256 | 45.7825 | 41 | P10C, P11S, P12S, P13C, P14C, P15C, P16C, P17C, P18C, P19S, P1C, P20S, P21S, P22S, P23S, P24S, P25S, P26S, P27S, P28C, P29C, P2C, P30S*, P32S*, P33C*, P34S*, P35S*, P36S*, P37S*, P38S*, P39S*, P3C, P40S*, P41S*, P42S, P4C, P5S, P6S, P7S, P8S, P9S |
| ASV2 | 796722 | 45.1851 | 38 | P11S, P12S, P13C, P14C, P15C, P16C, P17C, P18C, P19S, P1C, P20S, P21S, P22S, P23S, P24S, P25S, P26S, P27S, P29C, P2C, P30S*, P31C*, P32S*, P33C*, P34S*, P36S*, P37S*, P38S*, P39S*, P3C, P40S*, P41S*, P42S, P4C, P5S, P6S, P7S, P9S |
| ASV3 | 66152 | 3.7517 | 2 | P26S, P31C* |
| ASV4 | 36575 | 2.0743 | 1 | P11S |
| ASV5 | 22495 | 1.2758 | 2 | P12S, P16C |
| ASV6 | 9352 | 0.5304 | 1 | P35S* |
| ASV7 | 8869 | 0.503 | 1 | P14C |
| ASV8 | 7668 | 0.4349 | 3 | P14C, P26S, P31C* |
| ASV9 | 3913 | 0.2219 | 7 | P11S, P15C, P16C, P19S, P20S, P36S*, P39S* |
| ASV10 | 2128 | 0.1207 | 6 | P10C, P13C, P15C, P16C, P18C, P9S |
| ASV11 | 2111 | 0.1197 | 8 | P15C, P25S, P30S*, P32S*, P33C*, P34S*, P35S*, P3C |

**Supplementary Fig. 3.** Median joining network tree generated from (a) 11 ASVs of mt-ND1 and (b) 11 ASVs of mt-COX1. Each pie chart shows individual ASVs, and each population in the pie chart is indicated by a different color. The pie charts show the ASV distribution across all populations (insert tables). The number of mutations between ASVs is indicated with hatch marks along the connecting branches. The populations are grouped by county for AMOVA calculations using PopArt software.

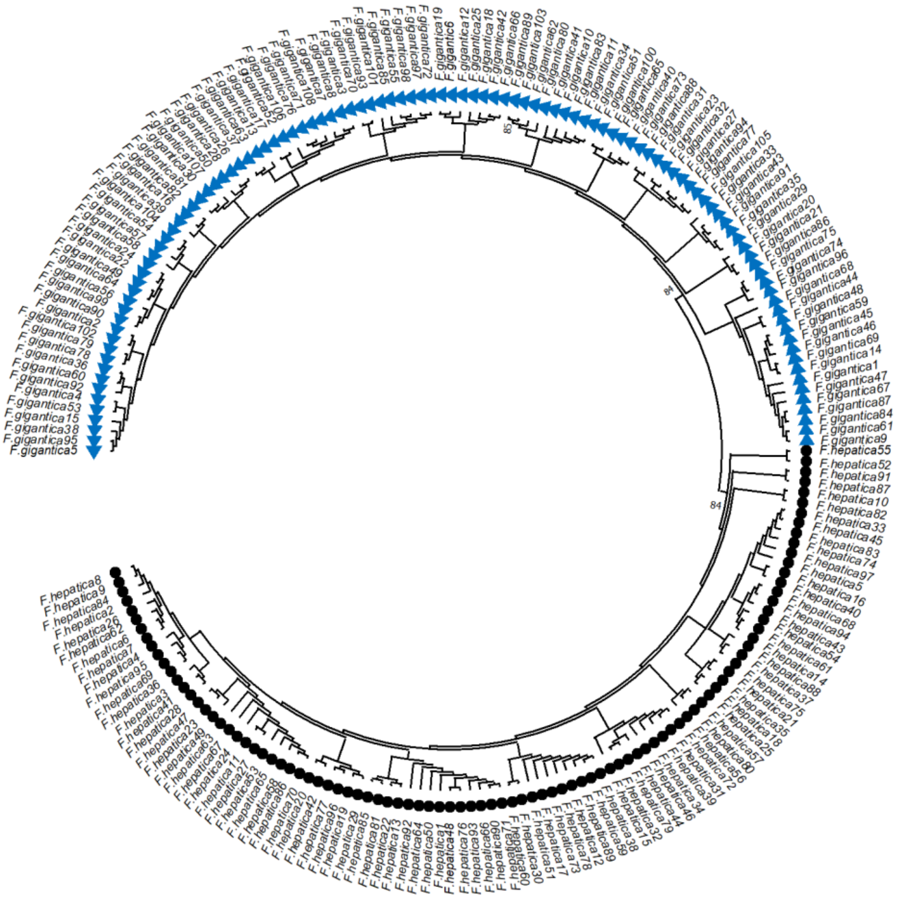

**Supplementary Fig. 4.** A Neighbour Joining tree for the *F. gigantica* and *F. hepatica* mt-COX1 loci was constructed using 2000 bootstrap replicates, from sequences identified via BLAST searches. A total of 205 sequences were included in the analysis, with 97 *F. hepatica* sequences marked by black circles and 108 *F. gigantica* sequences marked by blue triangles. The resulting tree showed clear clustering of the two species. *F. hepatica* validated reference sequences were then used to demultiplex mt-COX1 and classify reads generated from the Illumina MiSeq platform.
