## Supplementary Table 5 for "Limited genetic structure and high gene flow in *Fasciola hepatica* populations infecting ruminants in different geographic areas in the UK"

**Supplementary Table 5**. mt-COX1 primer sequences for amplifying cytochrome c oxidase subunit I region of *F. hepatica* mitochondrial DNA. The adaptor sequences are in italics, forward and reverse primers are underlined, and N's are random nucleotides added between Illumina's adaptor sequences and locus-specific primer. Modified phosphate bonds are added at positions indicated by the asterisks.

| Primer ID | Category/target region | Primer sequence | Product length | Remarks |
|---|---|---|---|---|
| mt-CO1 (i) 162 (For) | Cytochrome c oxidase subunit I | TGTTGTTACTGGGCATGGGA | 359bp | Forward primer |
| mt-CO1 (ii) 520 (Rev) | Cytochrome c oxidase subunit I | GACCCGTACCCTCATCCAAC | 359bp | Reverse primer |
| AD_For mt-CO1 (i) 162 | Cytochrome c oxidase subunit I | *TCGTCGGCAGCGTCAGATGTGTATAAGAGACAG*TGTTGTTACTGGGCATGG*G*A | - | Forward adapter primer |
| AD_For-1N mt-CO1 (i) 162 | Cytochrome c oxidase subunit I | *TCGTCGGCAGCGTCAGATGTGTATAAGAGACAG***N**TGTTGTTACTGGGCATGG*G*A | - | Forward adapter primer |
| AD_For-2N mt-CO1 (i) 162 | Cytochrome c oxidase subunit I | *TCGTCGGCAGCGTCAGATGTGTATAAGAGACAG***NN**TGTTGTTACTGGGCATGG*G*A | - | Forward adapter primer |
| AD_For-3N mt-CO1 (i) 162 | Cytochrome c oxidase subunit I | *TCGTCGGCAGCGTCAGATGTGTATAAGAGACAG***NNN**TGTTGTTACTGGGCATGG*G*A | - | Forward adapter primer |
| AD_Rev mt-CO1 (ii) 520 | Cytochrome c oxidase subunit I | *GTCTCGTGGGCTCGGAGATGTGTATAAGAGACAG*GACCCGTACCCTCATCCA*A*C | - | Reverse adapter primer |
| AD_Rev-1N mt-CO1 (ii) 520 | Cytochrome c oxidase subunit I | *GTCTCGTGGGCTCGGAGATGTGTATAAGAGACAG***N**GACCCGTACCCTCATCCA*A*C | - | Reverse adapter primer |
| AD_ Rev-2N mt-CO1 (ii) 520 | Cytochrome c oxidase subunit I | *GTCTCGTGGGCTCGGAGATGTGTATAAGAGACAG***NN**GACCCGTACCCTCATCCA*A*C | - | Reverse adapter primer |
| AD_ Rev-3N mt-CO1 (ii) 520 | Cytochrome c oxidase subunit I | *GTCTCGTGGGCTCGGAGATGTGTATAAGAGACAG***NNN**GACCCGTACCCTCATCCA*A*C | - | Reverse adapter primer |
